# The IQGAP1-Claudin Cell Junction Axis in Cisplatin-induced Kidney Damage

**DOI:** 10.1101/2024.12.21.629915

**Authors:** Yusuf Barudi, Xiuzhen Fan, Kenneth W. Jenkins, Shubhra Kanti Dey, Evgeny Yakirevich, Mahasin A Osman

## Abstract

Kidney damage resulting from nephrotoxicity is a major side effect of chemotherapy. Cisplatin is an effective antineoplastic agent broadly used in oncology where it interferes with DNA replication in dividing cells, however, its mechanism in kidney damage remains unclear. We tested the hypothesis that cisplatin displaces cell-contact proteins such as IQGAP1-claudin complex in the kidney tubules, causing epithelial cell dissociation and kidney damage. Employing a multifaceted approach, using mutant analyses and optimal cisplatin dose, this hypothesis was tested in cell culture and *iqgap1^−/-^* mouse models. Cisplatin inhibited cell proliferation and migration in an IQGAP1-dependent manner, displaced IQGAP1 from cell junctions and altered the expression level of key junctional markers, including claudins 2/4/8 and nephrin. Similar effects were observed in animal models where cisplatin treatment and loss of IQGAP1 had additive effects. These functional outcomes are accounted for by suppression of Akt1/PKB survival and ERK1/2 proliferation signals and activation of JNK-GSK3αβ stress and inflammatory signal, widely implicated in kidney injury. These findings present IQGAP1-claudin axis as biomarkers and therapeutic targets in kidney damage and pave the way for formulating new analogs that eliminate cisplatin adverse side effects while preserving its antineoplastic efficacy.

## Introduction

Epithelial cell adhesion is established and maintained by adheren and tight junction protein complexes that ensure epithelial cell polarity, create a tight seal of the lumen and serve as a platform for regulating protein trafficking and signaling.^1,2,3^ Dysfunction of any of the components of these complexes has been reported in a myriad of pathological states, ranging from cancer where the loss of cell polarity is the main culprit^4^ to bowel inflammatory diseases^5^ and polycystic kidney disease, where the loss of barrier function disrupts paracellular ion transport homeostasis.^6^

These adhesion complexes are organized by scaffolding proteins that often participate in the adhesion process.^2,7^ An example is the signaling scaffold IQGAP1, a key regulator of epithelial cell adhesion.^8,9^ Several studies showed that it controls renal epithelial cell polarity and adhesion both at the adheren and tight junctions to maintain the integrity and function of glomerular filtration *in vitro*.^9,10^ In the mouse, IQGAP1 localizes to nephron proximal and distal tubules,^11^ suggesting a role in secretion, but a role in kidney pathology remains undefined. Furthermore, we demonstrated that IQGAP1 modulates the integrity of tight junctions by interplaying claudin 2/4 expression and localization in the model Madin-Darby Canine kidney (MDCK) cells.^9^ The ramification of this modulation remained to be analyzed.

Claudins are a large family of transmembrane proteins essential for the formation and maintenance of epithelial tight junctions, acting both as paracellular pores and barriers.^12^ As such, in renal tubules, they determine permeability and selectivity of the various nephrons.^12^ The expression pattern of each claudin species varies considerably among the various tissues.^13^ Correlation between the expression level of claudin-3 and -4 with overall survival in clear cell renal cell carcinomas, suggested their potential as diagnostic and prognostic markers.^14^ Several studies demonstrated the utility of claudin 4 monoclonal antibodies in treating cancer or enhancing drug absorption by the paracellular permeation route.^15,16,17^ Claudins role in kidney function and pathology has been emerging.^18^

Kidney epithelial damage can occur in response to many diseases and therapeutic agents, such as the nephrotoxicity resulting from chemotherapy.^19,20^ Greater than 30% of chemotherapy-treated cancer patients develop renal injury despite the absence of other risk factors. ^20,21,22,23^ Many chemotherapeutic agents cause kidney-specific injury, including the platinum-based DNA synthesis modulator cisplatin, which plays a major role in oncology as a critical adjuvant, neoadjuvant and base standard of care anticancer management agent.^22,23,24,25,26^ Kidney damage is the major adverse side effect of cisplatin arising from apoptosis of tubular cells,^27,28,29,30^ however, its mechanism of action remains to be defined. Likewise, targeted cancer therapy also leads to kidney damage.^31^ Thus, determining the mechanisms underlying such injury is of crucial clinical and basic biological importance.

Here we tested the hypothesis that cisplatin impairs the IQGAP1-claudin junctional complex in kidney epithelia leading to dissociation of the cell-cell contacts and thus kidney damage. We employed a multifaceted approach, using mutant analyses and cisplatin to test this hypothesis in cell culture and in *iqgap1^−/-^* mouse model. The results show that IQGAP1 and claudin 4 colocalize at cell junctions in MDCK cells and in the distal tubules of mouse kidney. Cisplatin inhibited cell proliferation and migration in an IQGAP1-dependent manner, displaced IQGAP1-claudin 4 complex from cell junctions and altered downstream signaling pathways. Displacement of IQGAP1 appears to accompany differential expression of several other junctional markers. The insights gained from the results are twofold. First, they support the notion that cisplatin likely exerts an additional therapeutic effect besides tumor DNA damage by dissociating cell junctions in tumor tissues. Second, cisplatin inhibits cell migration involved in cancer cell invasion as well as in tissue damage repair, and thus by the same token causes kidney tubular damage. These functional outcomes are accounted for by activation of C-Jun N-terminal Kinase (JNK) stress signal, suppression of the serine/threonine kinases Akt/PKB and the member of the mitogen-activated kinases (MAPKs) extracellular signal-regulated kinase 1/2 (ERK1/2) and the consequential activation of the Akt1-substrate the glycogen synthase kinase 3 (GSK3αβ), widely implicated in kidney injury. These findings set the stage for better understanding of epithelial cell biology as well as for devising the next generations of effective analogs that eliminate cisplatin adverse side effects on the kidney while preserving its antineoplastic efficacy.

## Results

### Cisplatin Inhibits Renal Cell Proliferation, Disrupts Cell Contacts and Displaces IQGAP1-cluadin4

The hypothesis that cisplatin disrupts cell adhesion was tested in the model Madin-Darby Canine Kidney (MDCK) cell line we used previously to identify Claudin 2/4 as IQGAP1 partners in regulating tight junctions.^9^ Treatment with 10μM cisplatin overnight (one day) led to thickening of cell membranes and apparent dissociation of cell contacts (Fig. 1A, upper panels). By 48 hrs. the cell contacts were diminished and the cell nuclei enlarged (Figure 1A, lower panels). Next, the effect of the same cisplatin dose on cell proliferation was examined. Both in MDCK and human embryonic kidney (HEK293) cell lines, cisplatin significantly inhibited cell proliferation (Figs. 1B and C). To determine whether this effect occurs via IQGAP1, proliferation in cells expressing the different mutants of IQGAP1 was quantified. While cisplatin inhibited proliferation in control, it did more so in cells expressing IQGAP1 with more significant effect seen in cells expressing the dominant negative mutant IR-WW (W+cis, Fig1A). This result suggests that IQGAP1 overexpression exacerbates the cisplatin effect on cell proliferation. Next, we examined cisplatin effect on IQGAP1-claudin 4 localization. In DMSO-treated control cells, endogenous IQGAP1 and claudin 4 localized to cell-cell contacts and also were found in cytoplasmic puncta (Fig. 2B, upper panels). Treatment with 10μM cisplatin significantly reduced IQGAP1-claudin 4 localization in the cell contacts, but IQGAP1 aggregates and claudin 4 puncta remained in the cytoplasm (Fig. 2B, lower panels & C). Overall, cisplatin seems to inhibit cell proliferation in human and canine kidney cells likely via targeting IQGAP1-claudin 4 at cell contacts. It was unclear, however, whether cisplatin simply displaces IQGAP1-claudin 4 from cell contacts or reduces their expression, which we examined next.

**Figure 1.**
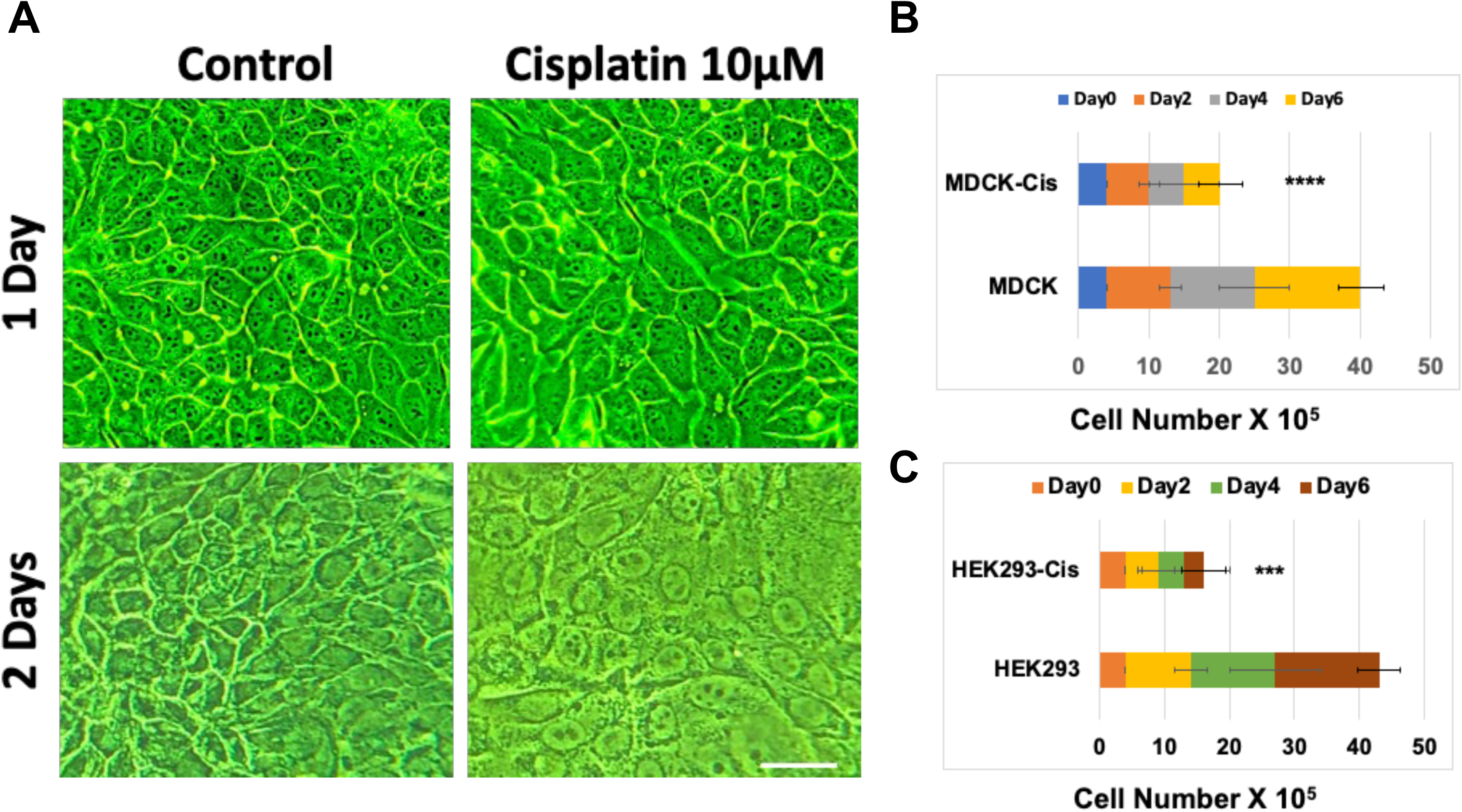
Cisplatin Inhibits Cell Proliferation and Disrupts Cell Contacts in Renal Cell lines. **A.** A widefield micrograph of live confluent MDCK cell lines that were treated with DMSO drug vehicle as control, left panels or with 10mM cisplatin, right panels. The cells were photographed after 1-day, upper panels and after 2 days, lower panels using brightfield microscopy. Scale bar 100 μm. **B.** Cell proliferation rates of control (MDCK) and 10μM cisplatin-treated (MDCK-cis) was compared in MDCK cell lines that were seeded at 10^5^ and counted every other day for 6 days to measure the saturation density with a cell counter. **C.** Cell proliferation rates of control (HEK293) and 10μM cisplatin-treated (HEK293-cis) was compared in human embryonic kidney HEK293 cell lines that were seeded at 10^5^ and counted every other day for 6 days to measure the saturation density with a cell counter.

**Figure 2.**
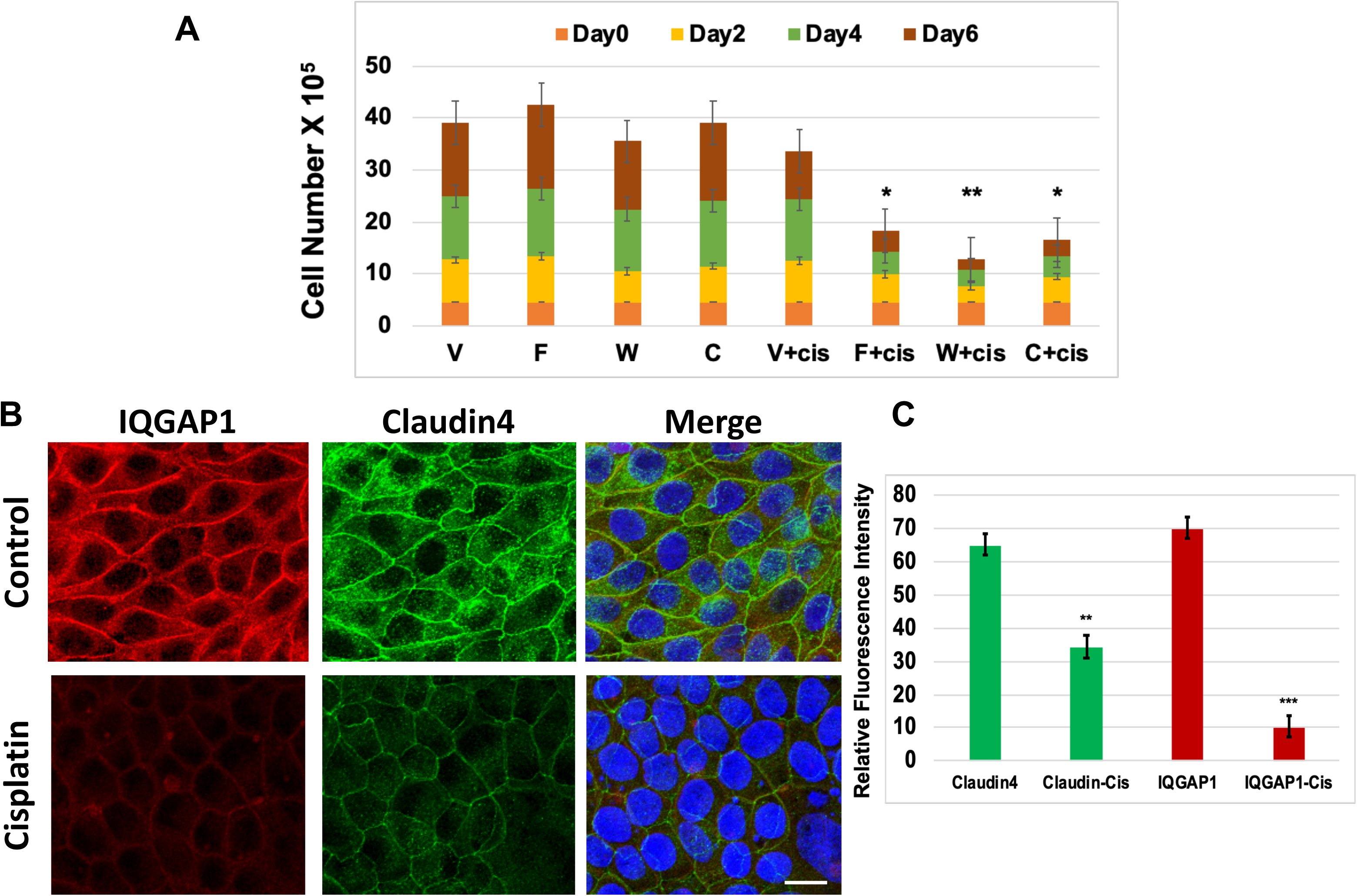
Cisplatin Displaces IQGAP1-claudin 4 complex from cell contacts. **A.** Cell proliferation rates of MDCK cell expressing the vector control (V) or full-length IQGAP1 (F), dominant negative IQGAP1-IR-WW (W) or dominant active IQGAP1-C that were treated with vehicle or 10μM cisplatin. The cells were seeded at the same number and counted every other day for 6 days to measure the cell saturation density with a cell counter. **B.** A confocal micrograph of confluent MDCK cell lines that were treated with DMSO drug vehicle as control, top panels or with 10mM cisplatin, lower panels and triple-stained with IQGAP1 antibodies (red), claudin 4 (green) and DAPI (purple for nuclei). C. Quantification of the signal intensities of Claudin 4, and IQGAP1 in control and treated cells. Scale bar 25μm. The error bars represent the mean ± standard deviation. * *p*< 0.05, ** *p<* 0.01, *** *p* < 0.001

### Cisplatin Differentially Alters the Expression Levels of Junctional Proteins and Activates JNK signal

Cisplatin effect on gene expression levels of several junctional proteins was examined both at the mRNA and protein levels. While the mRNA levels of claudin 2 and claudin 8 was significantly reduced in cisplatin treated MDCK cells compared to their controls, the mRNA levels of IQGAP1 and claudin 4 were significantly increased in cisplatin treated MDCK cells compared to their controls (Fig. 3A). The protein levels of the junctional complex markers did not always correlate with the mRNA level in response to cisplatin (Figs. 3A, 3B & C). Cisplatin treatment appears to significantly reduce claudin 4 and 2 protein level whereas that of claudin 8 and nephrin was increased (Fig 3B & C). The increase in IQGAP1 protein level in these cells appears to be statistically insignificant (Fig. 3B & C). Because IQGAP1 is a kinase scaffold the effect of cisplatin on expression and activity levels of select kinases associated with IQGAP1 pathway was examined. The levels of total JNK 46 and JNK 54 isoforms were significantly increased in response to cisplatin treatment (Figs. 3D & E). Notably, it was the levels of phosphorylated (active) JNK 46 monomers and JNK 54 dimers that was significantly increased in response to cisplatin (Figs. 3D & F). The functional outcome of these results was evaluated in terms of cell migration relative to IQGAP1 expression.

**Figure 3.**
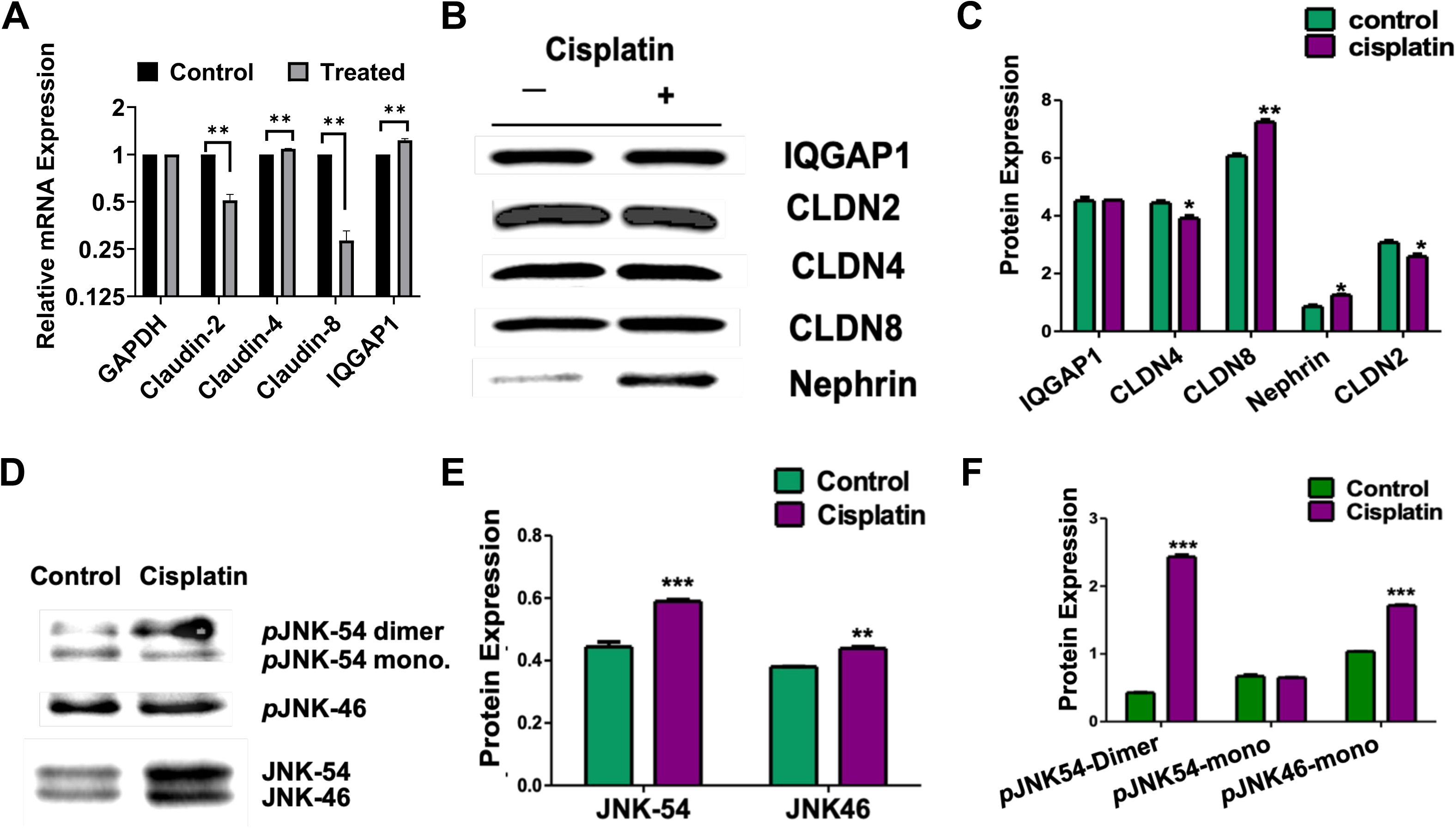
Cisplatin Differentially Alters the Expression Levels of Junctional Proteins and Activates JNK signal in MDCK cells. **A.** The mRNA levels of control (black bars) and cisplatin-treated (grey bars) MDCK cells was compared, using at least 3 reference genes, including the displayed GAPDH message (first two bars). **B.** A representative Western blot displaying the comparison between the protein expression levels of several known junctional protein markers. **C.** Quantification of the protein expression levels from blots of lysates isolated from control (green bars) and cisplatin-treated (purple bars) MDCK cells. **D.** Expression level of total and active Jun kinase (JNK) isoforms in control and treated MDCK cells. **E.** Quantification of total JNK isoforms and **F.** Quantification of the phospho-forms (active) JNK isoforms in the control and treated MDCK lysates. The error bars represent the mean ± standard deviation for n = 3 biological replica. * *p*< 0.05, ** *p<* 0.01, *** *p* < 0.001

### Cisplatin Inhibits Cell Migration and is Ameliorated by IQGAP1 Overexpression

Because IQGAP1 is a key regulator of migration in a variety of cell lines, and cell migration is crucial to organ healing relevant to kidney damage, we used wound-healing assays to evaluate the effects of 10μM cisplatin on cell migration. This effect was measured in MDCK cells (Fig 4A) and in wild type mouse embryonic fibroblasts (MEFs) overexpressing IQGAP1 full length (F), vector control (V), or MEFs harboring the *iqgap1* knockout (KO). Cisplatin inhibited cell migration in MDCK cells as compared to control up to 48 hr. post treatment (Fig. 4A & C). Similarly, in MEF control cells, cisplatin inhibited cell migration up to 48 hrs. post treatment (Fig. 4C, upper 2 panels). Overexpression of full length IQGAP1 was sufficient to overcome this effect, and the wound closed at 48 hrs. (Fig. 4C, middle 2 panels). In contrast, in cells harboring genetic knockout of *iqgap1*(KO), the wound was significantly wider, albeit a few cells reached the center of the wound (Figs. 4C, bottom 2 panels). Overall, these findings suggest that cisplatin inhibits cell migration and that IQGAP1overexpression can overcome cisplatin’s inhibition of cell migration. These finding were recapitulated preclinically in IQGAP1 animal model.

**Figure 4.**
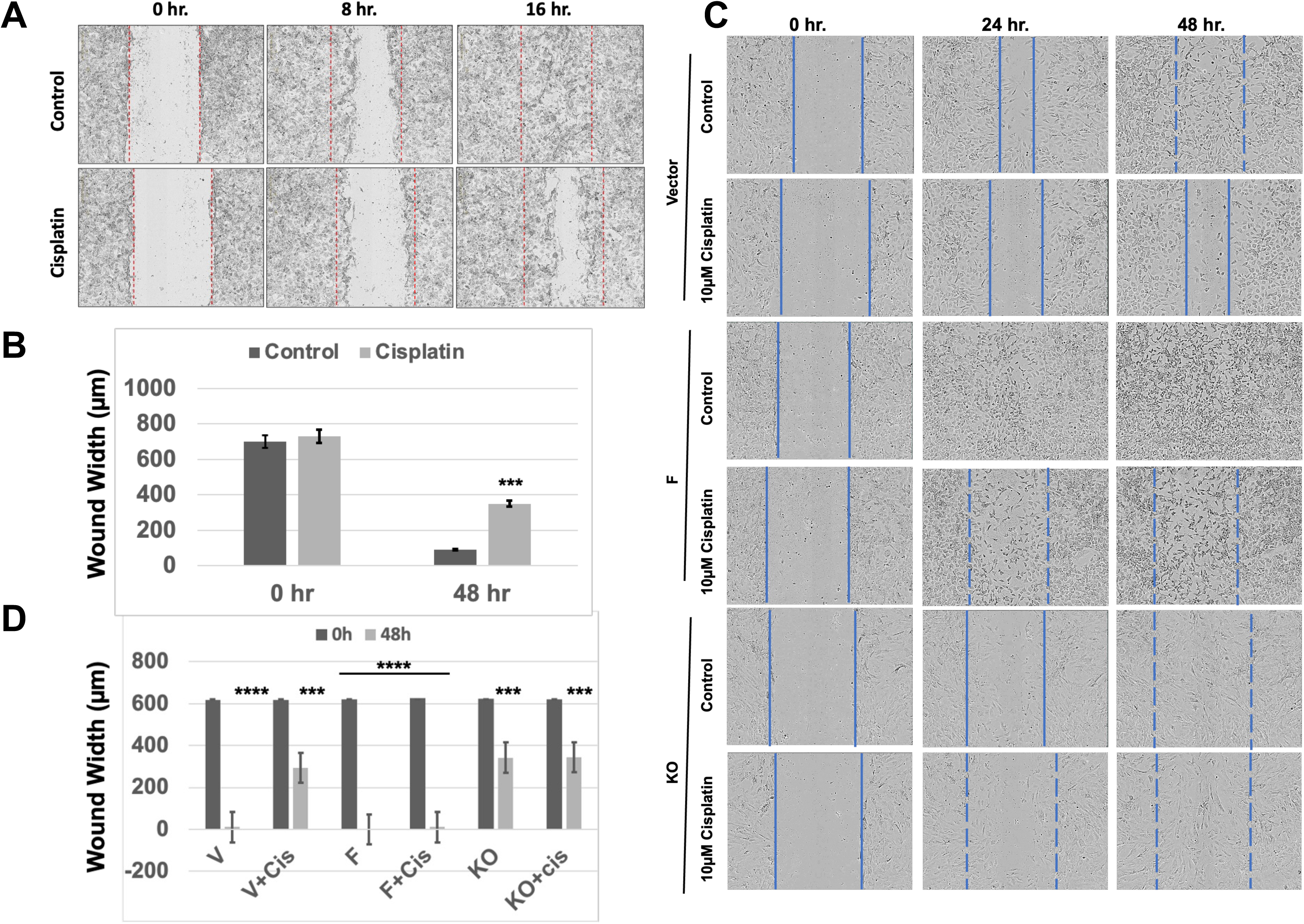
Cisplatin Inhibits Cell Migration and is Ameliorated by IQGAP1 Overexpression. **A.** Representative images of the effect of 10μM cisplatin on migration of control (upper panels) and treated (lower panels) MDCK cells. The cells were scratched and imaged up to 16 hr. post treatment. **B.** Quantification of the measurements of the wound width (migration rates) of the cells in Fig. 4A. The error bars are the mean ± S. D. for n = 3 biological replicas where each condition was run 8X. The results were considered significant at *** *p<* 0.001, **** *p* < 0.0001. **C.** Representative image of the effect of cisplatin on WT MEFs expressing the vector control, full-length IQGAP1 (F) or lacking *iqgap1* gene (KO). The cells were scratched and continuously imaged using the automated system incuCyte analyzer for 48 hrs. The solid lines represent the wound area whereas the dashed lines represent where the wound was. **D.** Quantification of the measurements of the wound width (migration rates) of the cells in Fig. 4C. The error bars are the mean ± S. D. for n = 3 biological replicas where each condition was run 8X. The results were considered significant compared to their respective control at *** *p<* 0.01, **** *p* < 0.0001.

### IQGAP1 Localizes to Tubules in Human Kidneys

Interactions of IQGAP1 and claudins in normal or diseased human kidney tissues has not been shown. Thus, we first investigated the distribution of IQGAP1 in normal human kidney samples to establish clinical relevance. IQGAP1 localized to distal tubules in medulla (Fig. 5A, left, S1), in the cortex at the brush border of distal tubules (Fig 5A middle) and at the proximal tubules around the glomerulus brush border (Fig. 5A, right). Previous reports showed that IQGAP1 localized to nephron distal convoluted tubules and moderately to proximal tubules.^11^ Furthermore, IQGAP1 localized to podocytes and glomerular endothelial cells in human nephrons.^32^ In podocytes, IQGAP1 localizes and interacts with the components of the slit diaphragm.^10^ These localization patterns resemble what we observed in mouse kidney and thus cisplatin effect was tested therein.

**Figure 5.**
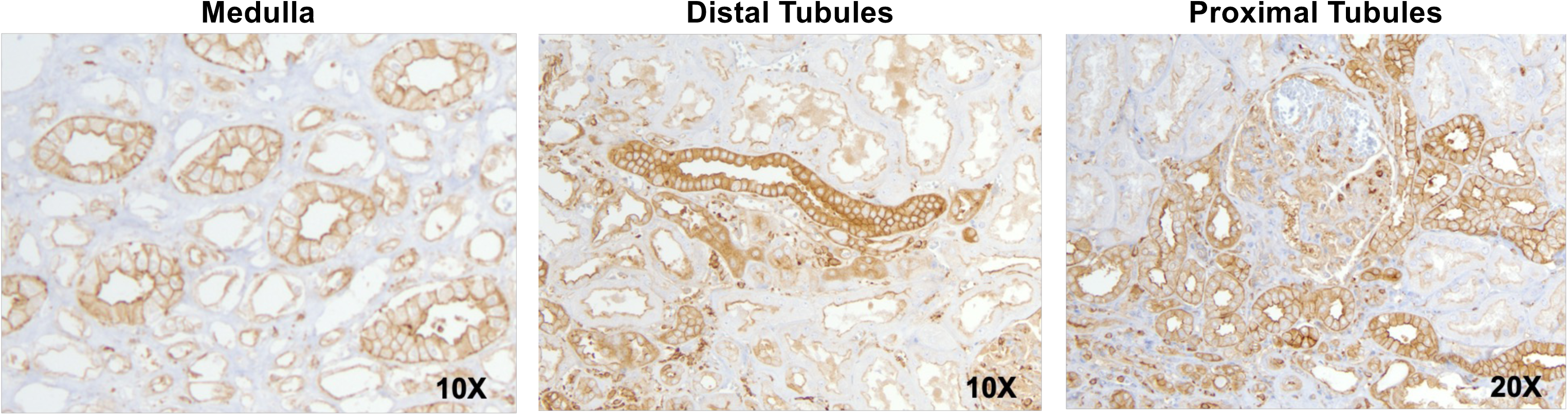
IQGAP1 Localizes to Tubules in Human Kidneys. **A.** Representative images of IQGAP1 localization in normal human kidney tissues. Left, IQGAP1 localizes to distal tubules in the medulla. Middle, IQGAP1 is found in the distal tubules in the cortex. Right, IQGAP1 is found in the proximal tubules around the glomerulus.

### Cisplatin Displaces IQGAP1 from Mouse Kidney Tubules

Two different immunohistochemical (IHC) staining methods were used to evaluate cisplatin effects on IQGAP1 localization in mouse kidneys. In wildtype control (WTC) kidneys stained with immunofluorescence, IQGAP1 localized to the cell contacts in the tubules and was found in the lumen/cytoplasm (Fig. 6A, left panel) and medulla (Fig. S1 A and B). In contrast, in the treated kidneys (WTT), IQGAP1 staining was obliterated from the tubules leaving a background level like the negative staining control (Fig.6A, middle and right panels). Due to the inherent autofluorescence in mouse tissues, we verified the results with chromogen IHC staining and it was clearer that cisplatin treatment displaced IQGAP1 from a more basal (cell contacts) structures in the wildtype control tubules (Fig. 6B, left panel, WTC) to a luminal/ cytoplasmic localization in the wildtype treated tissues (Fig. 6B, right panel, WTT). In addition to displacing IQGAP1, there was an overall increase in IQGAP1 staining in the medulla area (Fig, S1 A and B). Next, we examined the effect of IQGAP1 loss and cisplatin treatment on claudin 4 localization.

**Figure 6.**
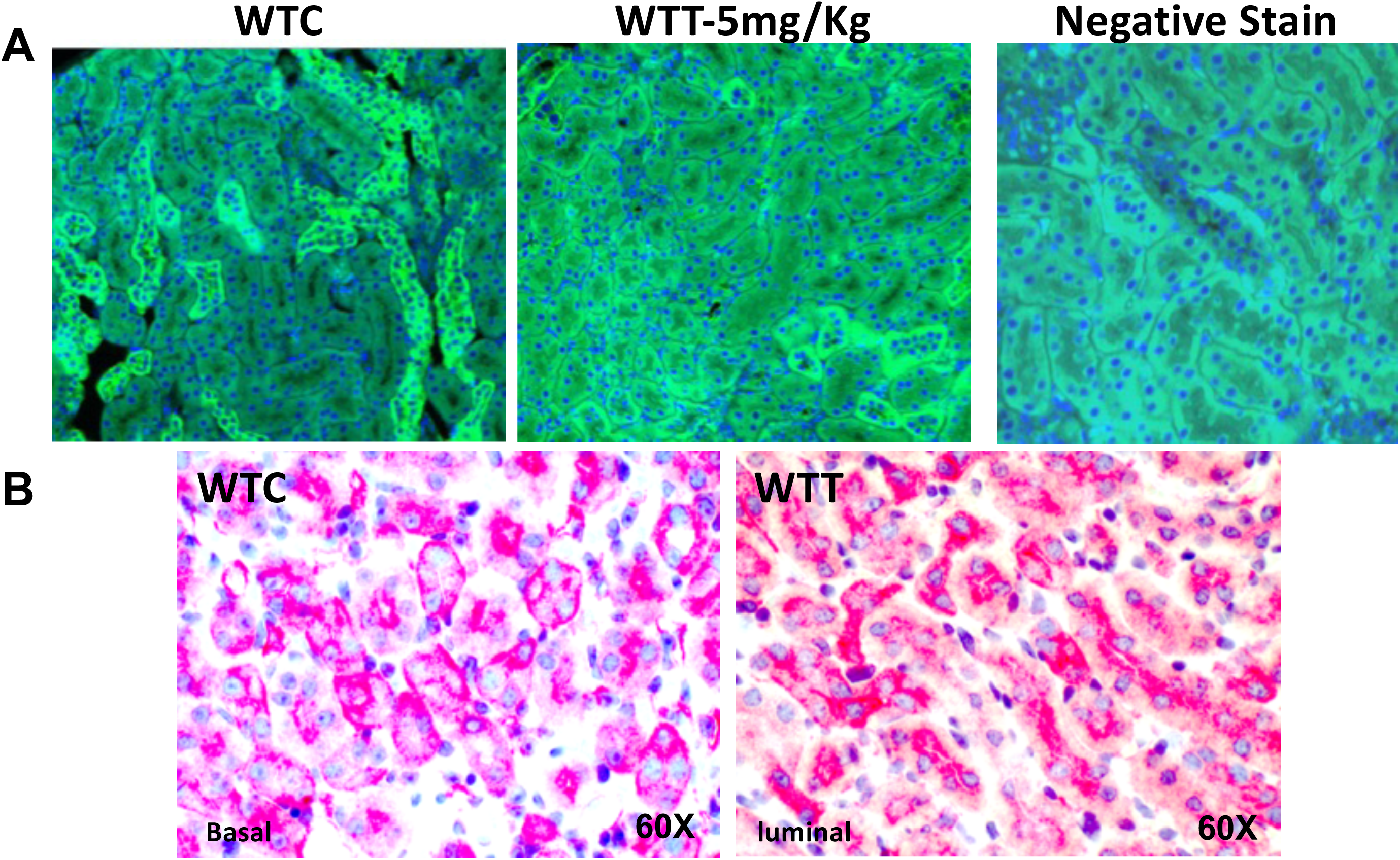
Cisplatin Displaces IQGAP1 from Mouse Kidney Tubules. **A.** Localization of IQGAP1 (green, Alexa Fluor 488) in wildtype control (WTC) kidney tissues (left panel), wildtype treated (WTT)with 5mg/Kg cisplatin (middle panel), the right panel represent a negative stain (IgG without IQGAP1 antibodies). **B.** Higher resolution (60X) images of WTC kidneys stained with chromogen, showing basal staining of IQGAP1 (left panel) and WTT kidney tissues showing luminal IQGAP1 staining.

### Loss of IQGAP1 and/or Cisplatin Treatment Enhance Claudin 4 Localization in Kidney Tubules

Previously, we showed that IQGAP1 silencing enhanced claudin 4 localization in the cell contacts.^9^ Here we compared IQGAP1-claudin 4 localization in kidneys isolated from wildtype and *iqgap1^−/-^* littermate controls or cisplatin treated. Fig. 7A shows that while cisplatin displaced IQGAP1 from the cell contacts in the tubules, it left claudin 4 correctly localized (Fig. 7A, lower panels). Additionally, genetic knockout in *iqgap1^−/-^*mice enhanced claudin 4 localization (Fig. 7B, upper panel) and expression level (Fig S1 C) and significantly more so when *iqgap1^−/-^*(KO) was combined with cisplatin treatment (Fig.7B, lower panels). Next, the effect of cisplatin on other known cell junctions markers was evaluated biochemically.

**Figure 7.**
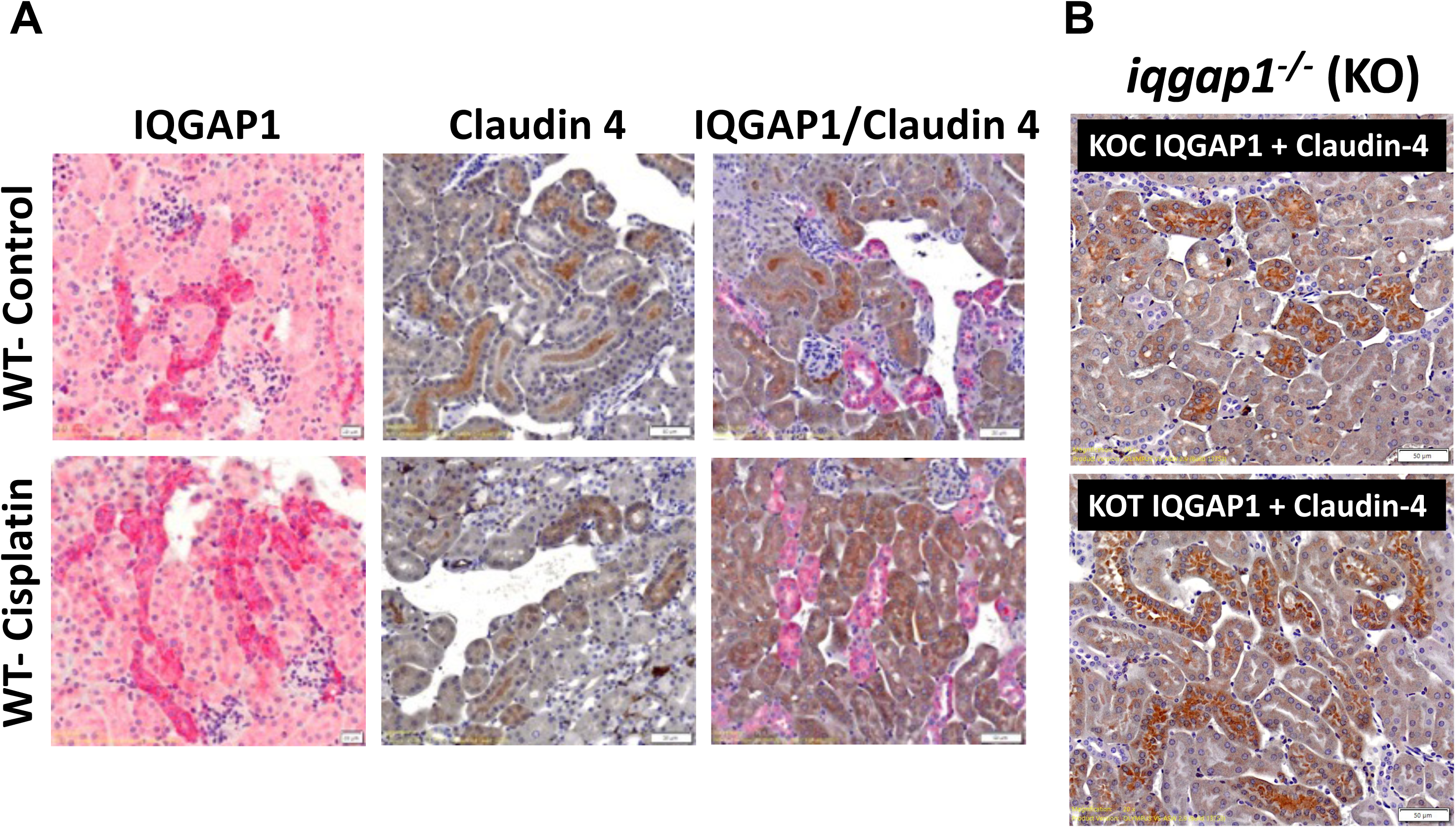
Loss of IQGAP1 and Cisplatin Treatment Enhance Claudin 4 Localization in Kidney Tubules. **A.** IQGAP1 and claudin staining in WT kidneys. Upper Panels, staining of IQGAP1 (pink), left panel and claudin 4 (brown, middle panel) and double stain (pink+ brown, right panel) in control WT kidney tissues. Lower panels. Same staining in cisplatin-treated kidney tissues. **B.** IQGAP1 and claudin staining in KO kidneys. Upper panel, representative double staining of IQGAP1 and claudin 4 in control KO kidney (KOC). Lower panel, double-staining of IQGAP1 and claudin 4 in cisplatin-treated KO kidney. Scale bar = 50 mm.

### Cisplatin Differentially Modulates the Expression Levels of Junctional Markers in Response to IQGAP1 Expression

The protein expression data from cell culture were recapitulated in wildtype (WT) and *iqgap1^−/-^* (KO) kidneys that were treated with cisplatin or the drug vehicle DMSO as control, using Western blot. In absence of IQGAP1 (KO), the expression levels of Nephrin, E-cadherin, and claudin 8 were significantly increased over that of WT (Fig. 8A&B). Upon cisplatin treatment, the levels of E-cadherin and claudin 8 were significantly decreased in the KO compared to WT (Fig. 8B). Interestingly, GAPDH protein level initially used as loading control was increased in the KO, but cisplatin treatment increased this level in WT compared to KO (Fig. 8B). Claudin 4 protein level did not significantly change with cisplatin application or IQGAP1 loss Fig 8A & B). Notably, cisplatin appears to significantly modulate claudin 2 level by enhancing it in the WT and reducing it in the KO. This was particularly apparent with claudin 2 dimers (50kD), and monomers (25kD) as seen in Fig. 8C. Overall, like in MDCK cell lines, cisplatin appears to influence the levels of the various components of the cell junction proteins differentially and in response to IQGAP1 loss (see the Discussion section). Next, we used a candidate approach to determine the signaling pathway involving the effect of cisplatin on the IQGAP1-junctional complex in the mouse model kidneys.

**Figure 8.**
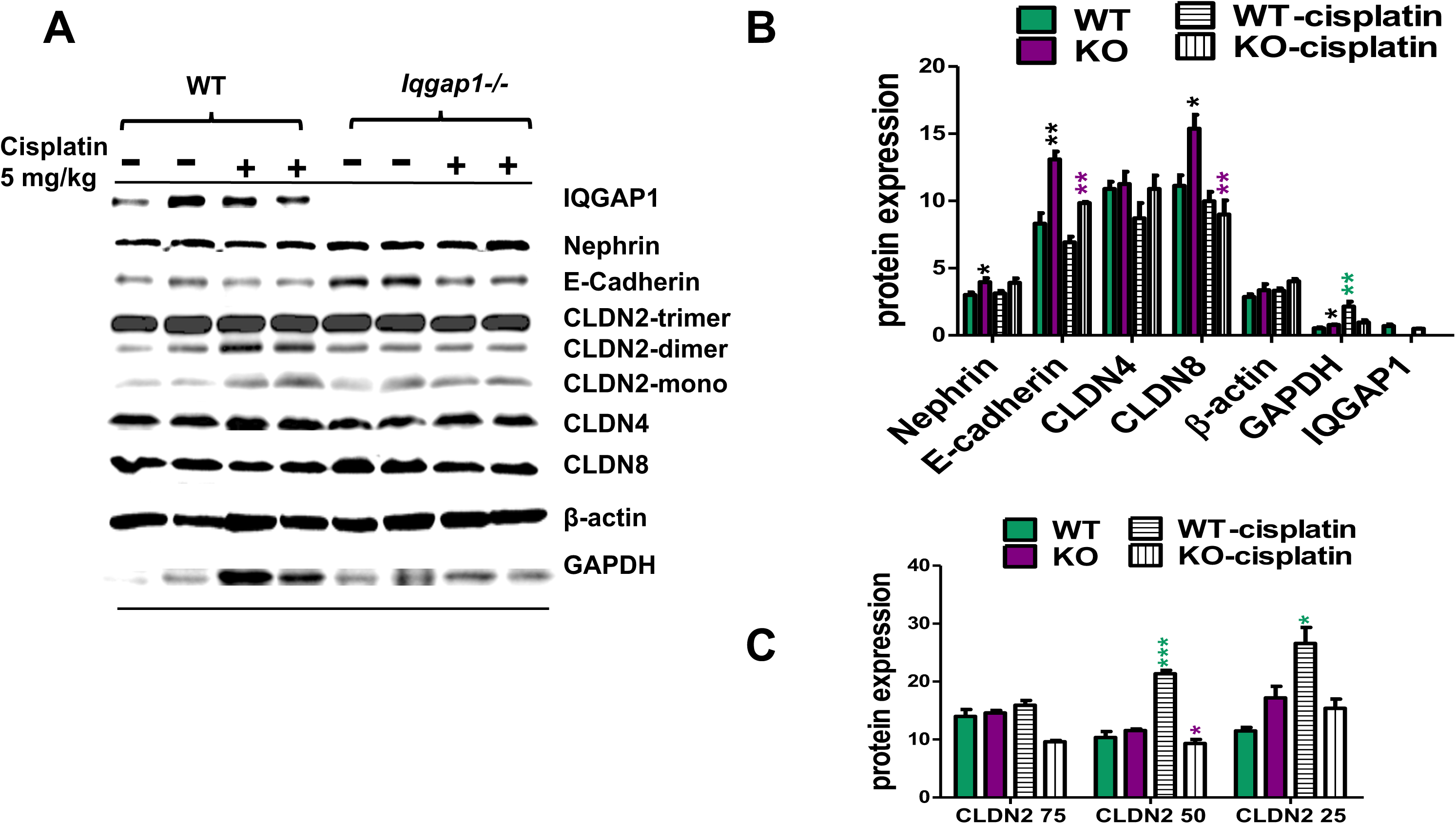
Cisplatin Differentially Modulates the Expression Levels of Junctional Markers in Response to IQGAP1 Expression. **A.** Comparison of various cell junction markers in WT and KO control and cisplatin-treated kidneys. A representative immunoblots of the indicated cell junction marker in control and treated WT and KO kidneys. **B**. Quantification of the expression levels (band intensities) of the markers, comparing treated to controls. **C.** Quantification of claudin 2 monomers and higher-level oligomers in control and treated WT and KO kidneys. The error bars are the mean ± S. D. for n = 5 biological replica with significance * *p*< 0.05, ** *p<* 0.01, *** *p* < 0.001 the color of the asterisk depicts the comparisons made: purple asterisk indicates KO treated compared to its control, green is WT treated compared to its control group.

### Cisplatin Modulates JNK, ERK1/2 and Akt1-GSK3ab Signaling in an IQGAP1-Dependent Manner

Extracts from wildtype and KO kidneys that were treated with cisplatin or vehicle control were quantified for the activity of known IQGAP1 signaling components (Fig. 9). Previously, we showed that silencing IQGAP1 activates JNK.^9^ Here we showed that in MDCK cells 10μM cisplatin increased the levels of total JNK and phosphorylated JNK 46 monomers and JNK 54 dimers significantly (Figs. 3D & F). The data in Fig. 9 A& B suggest that genetic loss of IQGAP1 increases the level of total JNK 54 while decreasing the level of total JNK 46 significantly (purple bars) over wild type (WT). Treatment of the mice with 5 mg/kg cisplatin led to a significant increase in total and *p*JNK 54 only in WT (Fig. 9 B, bars with horizontal stripes). By contrast, cisplatin led to a significant reduction in the levels of *p*JNK 46 and 54 in the KO as compared to the untreated controls (Fig. 9 B, bars with vertical stripes). Only total JNK 46 was increased in the KO in response to cisplatin. Together, these data suggest that cisplatin requires IQGAP1 to activate JNK.

**Figure 9.**
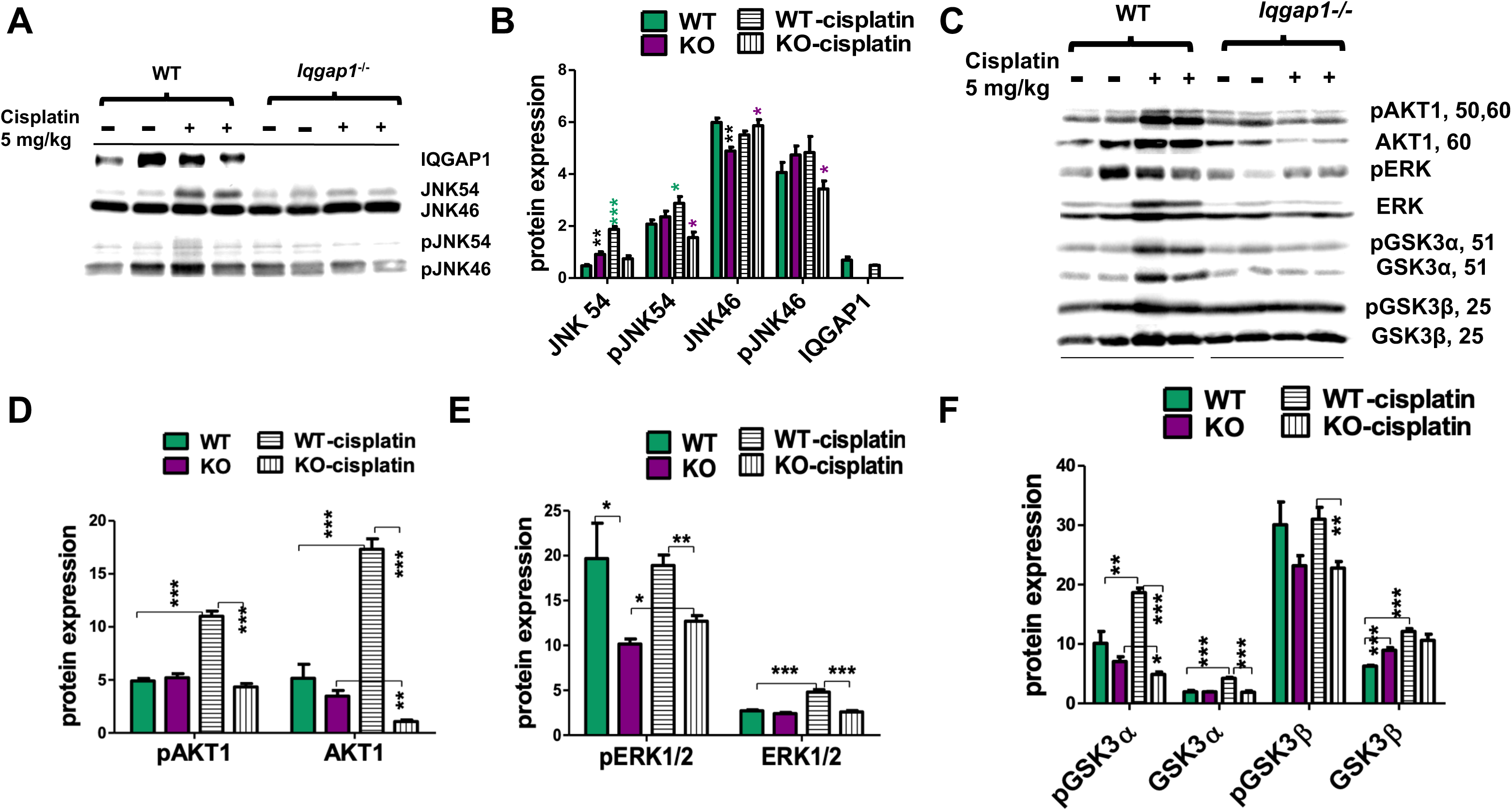
Cisplatin Interplays JNK, ERK1/2 and Akt1-GSK3αβ Signaling in an IQGAP1-Dependent Manner in Mice. **A.** a representative immunoblot for total and phospho-specific (active) JNK isoforms in kidney lysates from WT and KO (*iqgap1^−/-^*) mice. Lysates from two mice each WT and KO control (-) and treated (+) were selected at random for each blot (N =5) are presented in the blot. **B.** Quantification of the band intensities from 3 blots represented in 10A. The error bars are the mean ± S. D. for n = 3 biological replica * *p*< 0.05, ** *p<* 0.01, *** *p* < 0.001 comparing WT control groups to their treated groups (green asterisks) and KO control to their treated counterparts (purple asterisks). **C.** a representative immunoblot for total and phospho-specific (active) ERK1/2 and Akt1-GSK3αβ isoforms in kidney lysates from WT and KO (*iqgap1^−/-^*) mice. Lysates from two mice each WT and KO control (-) and treated (+) were selected at random for each blot (from N= 5) are presented in the blot. **D-F.** Quantification of the band intensities of the proteins presented in the blot in C. The error bars are the mean ± S. D. for n = 3 biological replica * *p*< 0.05, ** *p<* 0.01, *** *p* < 0.001, **** *p*< 0.0001. The comparisons are indicated with brackets.

Previously, we showed that IQGAP1 binds to and activates Akt1^34^ and thus we tested the effect of cisplatin on Akt expression and activity. Both total and *p*AKT^S473^ levels were increased significantly upon cisplatin treatment in the WT, but not in the KO (Fig 9C & D, bars with horizontal stripes). Thus, cisplatin application and loss of IQGAP1 appear to inhibit Akt1 activity. A similar pattern was observed for ERK1/2 expression level and activity with the exceptions that *p*ERK1/2 was higher in control WT compared to KO counterpart (Fig. 9E, green and purple bars), and an increase in *p*ERK1/2 was detected in the KO that was treated with cisplatin compared to its untreated control (Fig. 9E, compare bar with vertical lines and purple bar). The expression and activity (phosphorylated inhibition) of the Akt1 substrate GSK3αβ was consistent with the blunted Akt1 expression and activity (compare Fig. 9 D & F). The levels of *p*GSK3α were significantly lower in the cisplatin-treated KO compared to its control or to WT treated (compare Fig. 9F, bars with vertical stripes and purple). The expression level of total GSK3β was significantly higher in the untreated KO compared to WT (Fig. 9 F, compare green and purple bars), but cisplatin increased *p*GSK3β in wild type significantly.

### Methods Cell Culture

Madin-Darby Canine Kidney (MDCK) and human embryonic kidney 239 (HEK293) cells were purchased from ATCC and grown per the manufacturer’s instructions in DMEM containing 100-units/ml penicillin, 100 mg/ml streptomycin (Invitrogen) supplemented with 10% fetal bovine serum (FBS). The cells were maintained in a humidified incubator at 37°C and 5% CO_2_. The cells were discarded after 8 passages and replaced with an earlier passage from liquid nitrogen storage. The V5-IQGAP1 constructs, and the generation of stable cell lines was previously described.^33,34^ Mouse embryonic fibroblasts (MEFs) were isolated from wildtype and *iqgap1* knockout littermate mice and established as primary cells grown in DMEM supplied with calf serum and antibiotics. All chemicals (e.g. cisplatin, DMSO) were molecular biology grade obtained from Sigma-Aldrich or Fisher Scientific.

### Cell Proliferation Assay

Cell proliferation assays were compared both as growth in low serum (1%) or as the saturation density in high serum (10%) as previously described^33,34^ to evaluate proliferation rates of MDCK, HEK293, MEF cells or IQGAP1 mutants cell lines in response to cisplatin as compared to the cisplatin vehicle DMSO as control. Briefly, the cells were seeded at 4×10^4^ in multi-well plates and incubated overnight (16 hr.) to attach. The media were replaced with fresh media containing 10µM cisplatin or DMSO and changed with the respective media every other day for 6 days. At the indicated time points (2, 4, 6 days) the cells were washed with PBS, trypsinized and counted with a Countess II Automated Cell counter (Thermo Fisher Scientific Invitrogen).

### Fluorescence Immunocytochemistry (IF)

Cells were cultured in multiple-chamber slides (Nalge, Nunc), treated with cisplatin or DMSO vehicle as control for 48 hr. (2 days) and stained as previously.^35^ Briefly, the cells were washed with PBS and fixed in –20°C methanol for 10 min, permeabilized in PBS containing 0.1% Triton X-100 and blocked with 1% BSA in PBS, incubated at 4°C overnight with primary or control antibodies followed by secondary (Texas Red, Alexa Fluor 555, or Alexa Fluor 488, Molecular Probes) for 1 hr. at room temperature and the nuclei were counter-stained with DAPI (Sigma or Invitrogen) for 5 minutes at room temperature. After extensive washing with PBS and water, the slides were mounted and imaged with a Leica two-photon confocal microscope in our Integrated Facility. Image analyses were done using NIH ImageJ from at least 5 random fields for a total of ∼50 cells.

### Cell Migration Assays

The effect of cisplatin on cell migration was evaluated with the automated IncuCyte^©^ Live-Cell Analysis system (Agilent Technologies) in treated and control MDCK cells and in mouse embryonic fibroblasts (MEFs) isolated from WT and KO *iqgap1^−/-^* mice. The MDCK cells were seeded at 4×10^6^ in DMEM and allowed to grow to confluency. The monolayers were scratched using a Wound Maker^TM^ (Essen Bioscience) and washed 3X with PBS and treated with 10µM cisplatin or vehicle control. The cells were imaged every 6 hrs. for 48 hrs., using the IncuCyte Live-Cell Analysis System (Essen Bioscience). In comparing the MEFs, the variable growth rates of the cells was taken into account in seeding the cells in order to attain equal confluency (similar monolayers) as follows: In a 96-well plate, the wild type MEFs expressing the empty vector (V) were seeded at 40,000 cells per well in a 96-well plate, MEFs overexpressing full-length IQGAP1 (F) were seeded at 30,000 cells per well, and the knockout (KO) MEFs were seeded at 50,000 cells per well. The cells were cultured in DMEM media to 90% confluence, typically on the following day. A scratch was created in the center of the cell monolayer using the Wound Maker^TM^ (Essen Bioscience) followed by washing the plates three times with PBS to eliminate cell debris. The cell lines were then treated either with control (DMEM plus DMSO) or 10 μM cisplatin in DMEM media. Each condition was run in octuplicate (8X). Digital images were captured every 6 hours for 48 hours to track wound width accurately using the IncuCyte Live-Cell Analysis System. The images were imported from the IncuCyte, and Image J software was used to fine-tune brightness and contrast. The Images were analyzed, and the graph data were generated using the automated software algorithm functions from IncuCyte or Excel Microsoft software.

### Quantitative RNA Polymerase Reaction (qRT-PCR)

Log phase MDCK cells (80% confluency) were treated with 10 μM cisplatin or vehicle control for 48 hrs. The cells were processed using the Zymo Research Quick-RNA MicroPrep kit for total RNA extraction following the manufacturer instructions, and the RNA was quantified with a Nano drop (thermos-Fisher). The cDNA was generated using the iScript Reverse Transcriptase Supermix enzyme (Bio-Rad) and a Bio-Rad cycler. Primers for mouse or dog target genes were determined from the National Institute of Health GenBank and synthesized by Integrated DNA Technologies (IDT). Positive and negative controls alongside several reference genes, including those for TATA binding protein (TBP), Glyceraldehyde 3-phosphate dehydrogenase (GAPDH) and β-actin were used for qRT-PCR in 96-well plates in triplicates, using an Applied Biosystem machine. The *t*-test was calculated against each reference gene and *p* values equal or less than 0.05 were considered statistically significant. Human IQGAP1 primers have been described previously.^35^ The dog primers used for MDCK cells were as follows:

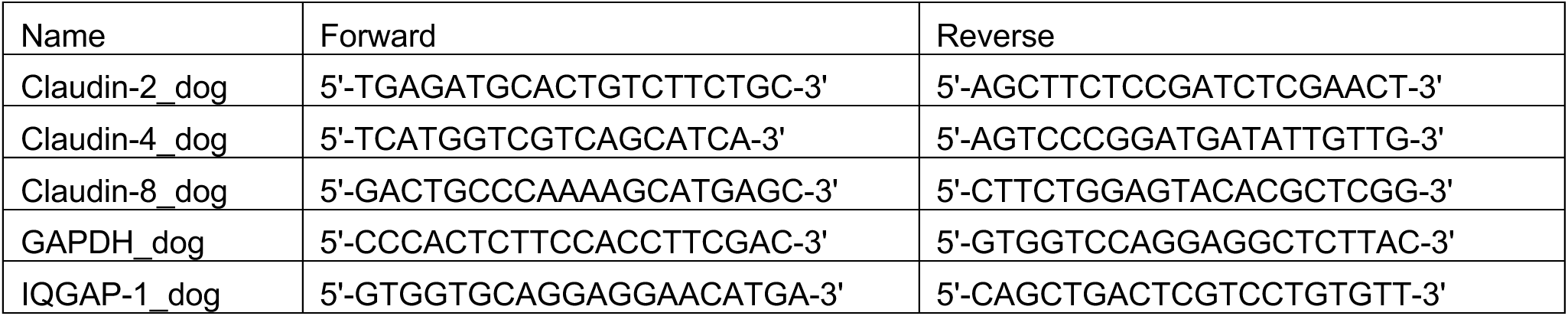

### Human Kidney Samples

Normal Kidney samples were obtained from Rohde Island Hospital of Brown University. Samples were collected with informed consent and the de-identified processed tissue slides were provided to the investigators in this study. Tissue samples were processed for IHC at the Rhode Island Hospital pathology facilities and annotated by the attending pathologist Dr. Evgeny Yakirevich.

### Animals and Study Approvals

Wild type and *iqgap1*^−/-^ (KO) mice (129/SVJ strain) were kindly provided from Dr. Sayeepriyadarshini Anakk at University of Illinois Urbana Champagne. The mice were bred in our pathogen free Department of Laboratory Research Animals (DLAR) facility, University of Toledo. Mice were fed a chow diet and maintained at 22 °C and 50% humidity with 12 hr. light/dark cycle. All animal procedures used in this study were approved by the Institutional Animal Care and Use Committee (IACUC) which is AAALAC and NIH accredited and comply with or exceed the NIH regulations. The animals were monitored daily by the DLAR staff, including a resident veterinarian and by the research staff. Eight weeks old littermate wild type or KO male mice were administered a single 5 mg/kg cisplatin (Sigma–Aldrich, St Louis, MO, USA) dose or phosphate buffer saline (PBS-DMSO vehicle) as a control via tail vein. Per power analyses, each group had 5 mice. After twelve weeks, the mice were sacrificed, and the harvested kidneys were split into halves and one half frozen in -80°C for biochemical studies and the other fixed in formalin overnight for histological studies.

### Fluorescence and Chromogen Immunohistochemistry (IHC)

Mice kidneys were fixed in 10% formaldehyde in PBS overnight, transferred to 70% ethanol before paraffin-embedded and 5-µm thick coronal sections were generated for IHC. After deparaffinization and hydration, the sections were incubated in 3% H_2_O_2_ solution in PBS at room temperature for 10 min. Antigen retrieval was performed in sodium citrate buffer (0.01 M, pH 6.0) in 95°C water bath for 20 minutes. Nonspecific antibody binding was blocked by incubation with 5% normal goat serum in PBS for 1 h at room temperature. Slides were stained overnight at 4 °C with the following primary antibodies: IQGAP1 (Santa Cruz, Cat. No. sc-1079, 1:200 dilution) and claudin 4 (1:200 dilution). The slides were subsequently washed and incubated with biotin-conjugated secondary antibodies for 30 min, and then with Horseradish Peroxidase Streptavidin (HRP Streptavidin) for 30 min. The sections were developed using the 3,3ʹ-diaminobenzidine (DAB) substrate for 5 minutes (SPlink HRP Rabbit Detection (DAB) kit, cat. no. D-03-18; OriGene Technologies, Inc.) and counterstained with hematoxylin QS (H-3404, Vector Laboratories). For IQGAP1-Claudin double-staining, IQGAP1 was stained red/magenta using ImmPACT Vector Red Substrate Kit, Alkaline Phosphatase. (SK-5105) and claudin was stained brown using Vectastain Elite ABC Universal kit, Peroxidase (PK-7200). The images were captured with an Olympus microscope.

### Western Blot (WB)

Lysis buffer [20 mmol/L Tris (pH 7.4), 137 mmol/L NaCl, 1% NP40, 10% glycerol supplemented with protease inhibitors, phosphatase inhibitors, 10 mM DTT and 5 mM ethylenediaminetetraacetic acid (EDTA)] was used to process cells and tissues. Control and treated samples were washed with ice cold tris buffered saline (TBS) and homogenized (cells were scraped or pipetted up and down 10 times with a 200 µl tip) into ice-cold lysis buffer on ice. The lysate was centrifuged at 1000 *g* for 20 minutes at 4°C to remove the cell debris. The supernatant was transferred to a fresh tube and protein concentration measured using the BioRad BCA kit. Total extracts 20μg (cell extracts) or 25μg (tissue extracts) were boiled for 5 min and loaded onto 10% or 15% SDS-Tris-glycine gel and resolved at room temperature.

Proteins were then transferred to PVDF membrane blocked in TBST blocking buffer (TBS pH 7.4, 0.1% Tween 20) and 5% nonfat milk (or BSA for phospho-proteins) for 1 hr. The membranes were probed with target-specific primary antibodies in TBST with 5% nonfat milk (or BSA for phosphor-protein) at room temperature for 2 hrs. or 4°C overnight. The membranes were washed in TBST 3X for 15 min and incubated with horseradish peroxidase (HRP)-conjugated appropriate secondary antibodies (1:7000, goat to mouse or rabbit) in TBST with 5% nonfat milk (or BSA for phosphor-protein) for one hr. followed by washing. The bands were visualized with ECL system and imaged by ChemiDoc XRS (Bio-Rad ) then analyzed by Image Lab software (version 6.0, Bio-Rad). We use Coomassie Brilliant Blue G-250 stained total protein loading on PVDF membrane as reference for quantifying the target protein^36^ followed by quantification with Image Lab Software. Primary antibodies dilution details are shown below.

### Antibodies

All antibodies were obtained from reliable biotech companies and validated in our lab by recognition of control vs. knockdown proteins with RNAi, tagged expressed proteins or extracts from *iqgap1* knockout mouse. Monoclonal antibodies for IQGAP1 rose in rabbit or mouse were previously described^34^ and were obtained from Pierce (Thermo-Fisher Scientific) or Santa Cruz Biotechnology (Santa Cruz, CA, Dallas, TX). Antibodies for claudins, nephrin, E-cadherin were from Santa Cruz Biotechnology and/or ABclonal Technology (Woburn, MA). Total and phospho-specific antibodies were purchased from Cell Signaling Technology (Beverly, MA), Fisher or Santa Cruz.

| Ab | Dilution | Ab | Dilution | Ab | Dilution |
| --- | --- | --- | --- | --- | --- |
| IQGAP1 (C-9) | 2000x | Claudin 2 | 1000x | Claudin 4 | 1500x |
| Claudin 8 | 2500x | Nephrin | 2000x | E-Cadherin | 2000x |
| GAPDH | 2000x | SAPK/JNK | 700x | pSAPK/JNK(Thr183/Tyr185) | 700x |
| pERK1/2 (44/42) | 700x | ERK 1/2 (C-9) | 2000x | GSK3$\alpha\beta$ | 1000x |
| AKT1 | 1000x | pAKT1 | 500x | pGSK3$\alpha\beta$ | 1500x |

### Statistical Analyses

All experiments were done in biological triplicates. Graph Pad Prism software (GraphPad Software, Inc., La Jolla, CA, USA) was used to plot the mean and standard errors, and statistical significance was determined by two-sided Student’s *t*-test for comparing specific protein band intensities of WB, or qPCR Ct values to related experiment groups. Statistical significance was set at *p* values equal to or less than 0.05. Fluorescence was quantified from 50 cells selected at random. IHC staining was quantified by blinded scorers, using whole organ staining from 10 tissue fields selected at random from the relevant areas, using the free online NIH-ImageJ.

## Discussion

Dysfunction of epithelial cell contact complexes associates with various pathological conditions, including renal cancer, bowel inflammatory and polycystic kidney diseases where their loss of function in cell polarity or disruption of paracellular ion transport have been implicated.^4,5,6^ Furthermore, systemic or targeted anticancer therapies lead to kidney damage.^19,20,31^ Unravelling the basic mechanisms of these dysfunctions is subject of active research. The results of this study support the hypothesis that the chemotherapeutic agent cisplatin displaces IQGAP1 and alters the expression and spatial distribution of its junctional partners causing epithelial cell dissociation, and thus likely contribute to kidney damage seen in chemotherapy treated patients. While the adheren junctions are thought to mediate cell-cell adhesion and the tight junctions regulate the ion and small molecule passage between cells, mounting evidence supports that the two are physically linked by scaffolding and signaling molecules.^37,38^ This notion has been enforced by the finding that the signaling scaffold IQGAP1 modulates both the adheren and the tight junction complexes and physically interacts with their components.^8,9^

Cisplatin inhibited cell proliferation in the kidney model MDCK and human kidney HEK293 cell lines (Fig. 1), which is consistent with its role in interfering with DNA replication in actively replicating cells^20,24^ such as in growing cell culture. Notably, expression of IQGAP1, which is oncogenic, appears to exacerbate cisplatin inhibition of cell proliferation, suggesting that cisplatin likely targets IQGAP1. An alternative explanation would be that expression of IQGAP1, a known oncoprotein, increases cell proliferation, which highlights the cisplatin effect on DNA replication, but does not exclude IQGAP1 targeting. In support of this notion, dissociation of cell contacts in cisplatin-treated MDCK cells was accompanied by specific displacement of IQGAP1-claudin 4 from these contacts (Fig. 2). This effect also was manifested at the protein expression levels where a significant reduction in claudin 4 was observed, accompanied by likely compensatory increases in IQGAP1 and claudin 4 mRNA levels (Fig. 3). This outcome was not uniform for all the junctional markers, hinting to the dynamic nature of these complexes. For example, claudin 8 mRNA level was reduced whereas its protein level increased (Fig. 3). While it is unclear at present how cisplatin affects gene expression at this low dose in the short duration of 2 days, it has been shown that cisplatin induces epigenetic and/or genetic alterations on numerous genes in cisplatin-resistant cells continuously exposed to cisplatin both *in vitro* and *in vivo*.^39^ Altered expression levels of claudin 4 has been reported in many types of cancers ranging from esophageal to prostate.^40^ In Crohn’s disease, upregulation of pore-forming claudin 2 and downregulation and redistribution of sealing claudins such as claudin 8 resulted in altered tight junction structure.^41,42^ In the kidney, claudins play a crucial role as pores and barriers, and as such they regulate the permeability and selectivity of various nephron segments in renal tubules.^12^ Consequently, some claudins have been implicated in genetic forms of hypertension and solute transport-related diseases.^12,43^ It is plausible that cisplatin-mediated displacement of IQGAP1 combined with altered expression levels of the different junction protein partners lead to alteration in the delicate balance/stoichiometry of the complexes, and thus compromising junctional integrity. In turn, this imbalance likely triggers a cell death signal emanating from the activation of JNK observed in cisplatin treated cells (Fig. 3). JNK is a known stress signal that serves as a hub in many cellular pathways leading to cell death,^44,45^ which may explain the loss of renal cells reported in kidney damage. Interestingly, cisplatin appears to induce activation of JNK 46 monomers and JNK 54 dimers. Dimerization plays a role in JNK activity.^46^ While the significance of this differential dimerization of the JNK subunits is unknown at present, it may generate a substrate selectivity to specify an apoptotic pathway activity under these conditions.

We surmised that cisplatin-mediated dissociation of cell contacts would enhance cell migration at least in cell culture. Instead, cisplatin inhibited cell migration both in MDCK cells and in wild type MEFs for up to 48 hr. post treatment (Fig.4). Interestingly, genetic knockout of IQGAP1 exacerbated the cisplatin inhibition of cell migration whereas overexpression of IQGAP1-F (full length) appears to abrogate the cisplatin inhibition (Fig. 4), again supporting the idea that cisplatin acts through IQGAP1. Mechanistically, this finding is consistent with the report that cisplatin generates cell stiffness via actin accumulation, which may inhibit cell migration.^47^ Given that IQGAP1 has a major role in actin dynamics,^48,49,50^ these findings again support the notion that cisplatin acts through IQGAP1.

To gain insights into the clinical significance of these observations, we first established that IQGAP1 localized to renal tubules in normal human kidney (Fig. 5), a known site of claudin action.^12^ IQGAP1 also localized to the medulla, the significance of which is to be investigated in the future. A similar localization pattern was observed in the mouse, thus lending credence to evaluating the effects of cisplatin on IQGAP1-junctional complexes in wild type and *iqgap1^−/-^* mouse model (Fig. 6). Like in human kidneys, IQGAP1 localized to tubules in wildtype cortex and cisplatin displaced IQGAP1 from these tubules (Fig. 6). However, there was an overall increase of IQGAP1 expression levels particularly in the medulla (Fig. S1), perhaps supporting an effect of cisplatin on IQGAP1 gene expression. Furthermore, there was an additive effect of *iqgap1* gene loss and cisplatin treatments. Loss of IQGAP1 led to an overall increase in claudin 4 level in whole kidney (Fig. S1), and cisplatin led to claudin 4 stabilization at the cell contacts and more so when the two conditions were combined (Fig. 7). These finding differ from those in MDCK cell (Fig. 3), where claudin 4 level and localization were reduced under such conditions likely due to sampling, physiological differences in cell culture and whole animal or species differences between dog and mouse. Regardless these data underscore the disruption in claudin 4 balance by cisplatin application and IQGAP1 loss. Claudin 4, forms pores with broader selectivity and thus causes an overall decrease in ion conductivity,^51^ therefore, it is intuitively possible that imbalanced claudin 4 level or chronic stabilization may contribute to kidney damage. While this assumption requires physiological studies in these models, cisplatin also altered the expression levels of other junctional proteins differentially and in an IQGAP1-dependent manner (Fig. 8). For instance, whereas the expression levels of Nephrin, E-cadherin, and claudin 8 was significantly increased in absence of IQGAP1, cisplatin treatment reversed this pattern (Fig. 9). By contrast, cisplatin enhanced claudin 2 level in wildtype and reduced it in the KO kidneys in agreement with our previous finding that silencing IQGAP1 in MDCK cells reduced expression level.^9^ Claudin 2 provides pore selectivity and increases conductivity at cell contacts (Amasheh et al., 2002; Furuse et al., 2001),^52,53^ and its activity has been shown to be regulated by dimerization^54^ as observed in Fig. 8. While the significance of the dimers is yet to be deciphered, overall, it appears that cisplatin working through IQGAP1 disrupts the dynamic balance between claudins 2/4 as well as other junctional protein partners of IQGAP1 thereby potentially contributing to eventual kidney damage.

Cisplatin action on IQGAP1 signaling components in mouse kidneys agree with the cell culture results albeit with some variance. While cisplatin modulated JNK signals, there were nuances in comparison to cell culture likely due to differences in physiological context. For instance, cisplatin significantly activated *p*JNK 54 only in WT and reduced the level of both *p*JNK 46 and 54 in the KO (Fig. 9). Taken together these novel findings support that IQGAP1 modulates JNK activity, and that cisplatin requires IQGAP1 to elicit JNK-stress signal. A similar scenario was observed for the known kinase effectors of IQGAP1; Akt1, ERK1/2 and GSK3αβ.^34^ Cisplatin did not enhance the levels of total Akt1 or active *p*AKT^S473^ and ERK1/2 in the absence of IQGAP1(Fig. 9). This finding is consistent with the report that PI3K/Akt1 signaling is activated by reduced claudin 4 expression,^55^ which occurs in cisplatin treated cells (Fig. 3). Thus, the combined effect of IQGAP1 genetic loss and increased expression or stabilization of claudin 4 appears to be an important factor in suppressing the Akt1 cell survival signal. Consequently, the Akt1-substrate GSK3α/β was activated by reduced serine-phosphorylation (Fig. 9). GSK3α/β whose action is regulated, at least in part, by an inhibitory serine-phosphorylation,^56^ is a known signaling node involved in acute kidney injury leading to chronic kidney disease.^57,58^ Work is underway to determine the receptor-mediated signal transduction events inputting into active JNK-GSK3α/β axis and the effectors mediating their downstream actions under these conditions.

In summary, we propose a model in which cisplatin, working through IQGAP1, disrupts the delicate structural and functional balance of cell junction proteins leading to the induction of JNK-GSK3α/β stress signal that likely compromise kidney epithelial ion and secretion balances, as well as cellular integrity, which underlie the observed cisplatin-associated kidney damage (Fig. 10 ). Based on these findings, future studies should include chemical synthesis of cisplatin analogs that preserve or enhance its efficacy in oncology on DNA replication while alleviating its deleterious effects on normal epithelial cell junctions.

**Figure 10.**
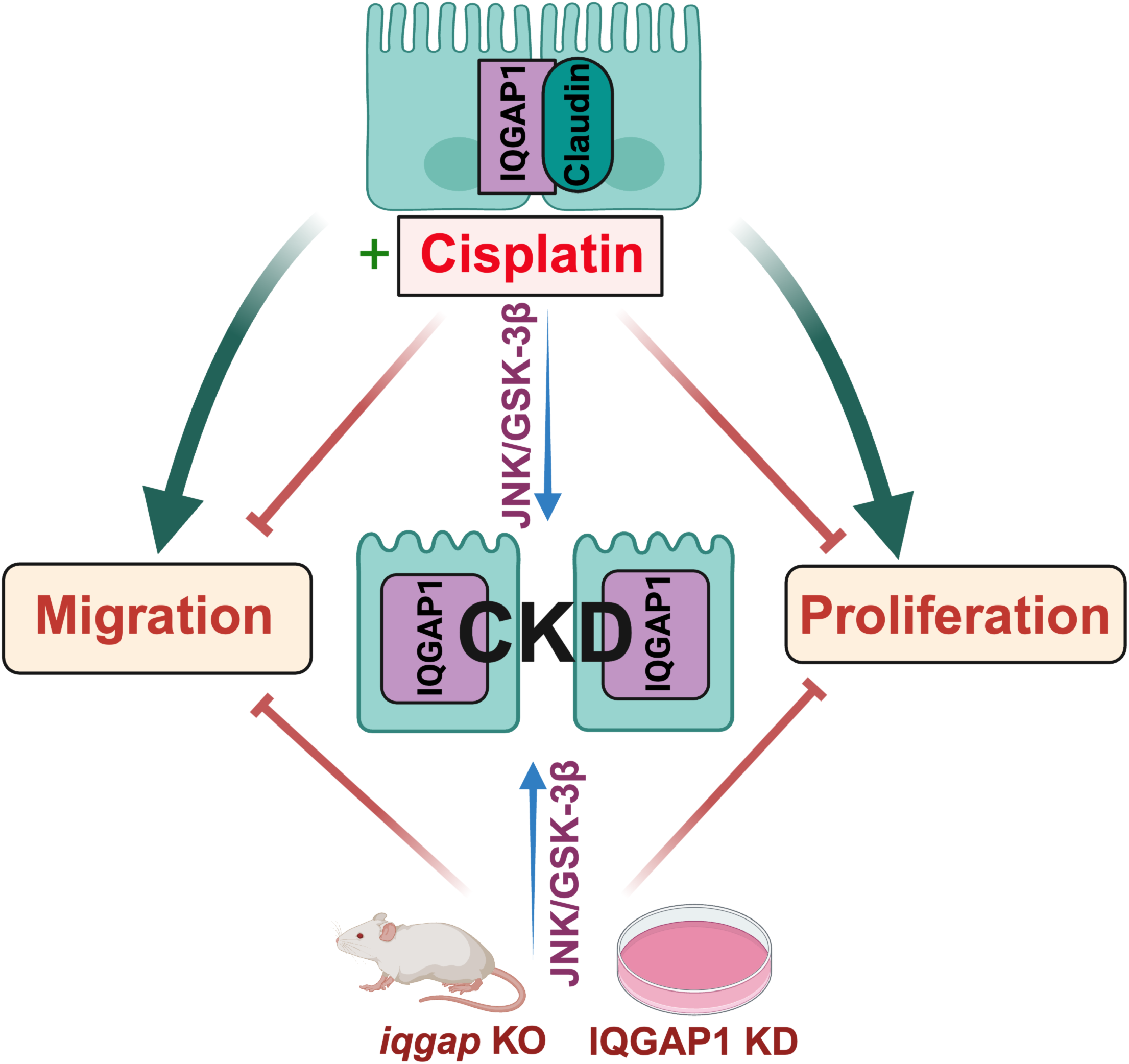
Model of cisplatin and/or IQGAP1 loss on cell junction proteins potentially leading to kidney damage. Upper image, normal kidney epithelia are polarized and connected with IQGAP1 scaffold and claudins as representative of intact junctional proteins. These cells can proliferate and migrate for wound healing. Middle image, cisplatin inhibits cell proliferation and migration, dissociates IQGAP1 from the cell contacts into the cytoplasm, and induces JNK-GSK3αβ stress signal thereby disrupting cell junctions, and potentially leading to chronic kidney damage (CKD). Lower image, loss of IQGAP1 via knockdown (KD) ^9^ or dominant negative mutants,^33^ or genetic knockout (KO) also leads to epithelial cells dissociation and induction of JNK/ GSK3αβ stress signal. The effects of cisplatin and IQGAP1 loss is additive on the loss of epithelial polarity and cell contacts.

## Data Sharing Statement

Details for primer sequences, antibodies and all relevant data for this study are included within the manuscript. Readers may contact the corresponding author with any reasonable requests.

## Acknowledgements

This study was supported in part by deAece-Koch Memorial Endowment seed fund and startup fund to MO from the University of Toledo. Former members of the Osman Lab Dr. James Antonisammy, Rawan Moussa, and Meghana Reddy Medini contributed to the earlier stages of this study. Varun Iyer helped with editing and verifying the references. Christopher Wojciechowski and Mahmood Meqdad helped with graphical art drawing, using BioRender software. We appreciate Mr. Allan Schroering of the UToledo Integrated Facilities for help with IHC tissue processing. We thank Ms. Kara Lombardo, Senior Research Assistant and Ms. Ashlee Sturtevant, the Department of Pathology, the Rhode Island Hospital of Brown University for expert assistance with chromogen IHC of human samples.

## Author Contribution

Y.B., X.F. K.J. S. D. K and E. Y performed experiments and edited the manuscript. M.A. O. designed the study, performed experiments, drafted and edited the manuscript.

---

**Figure S1.**
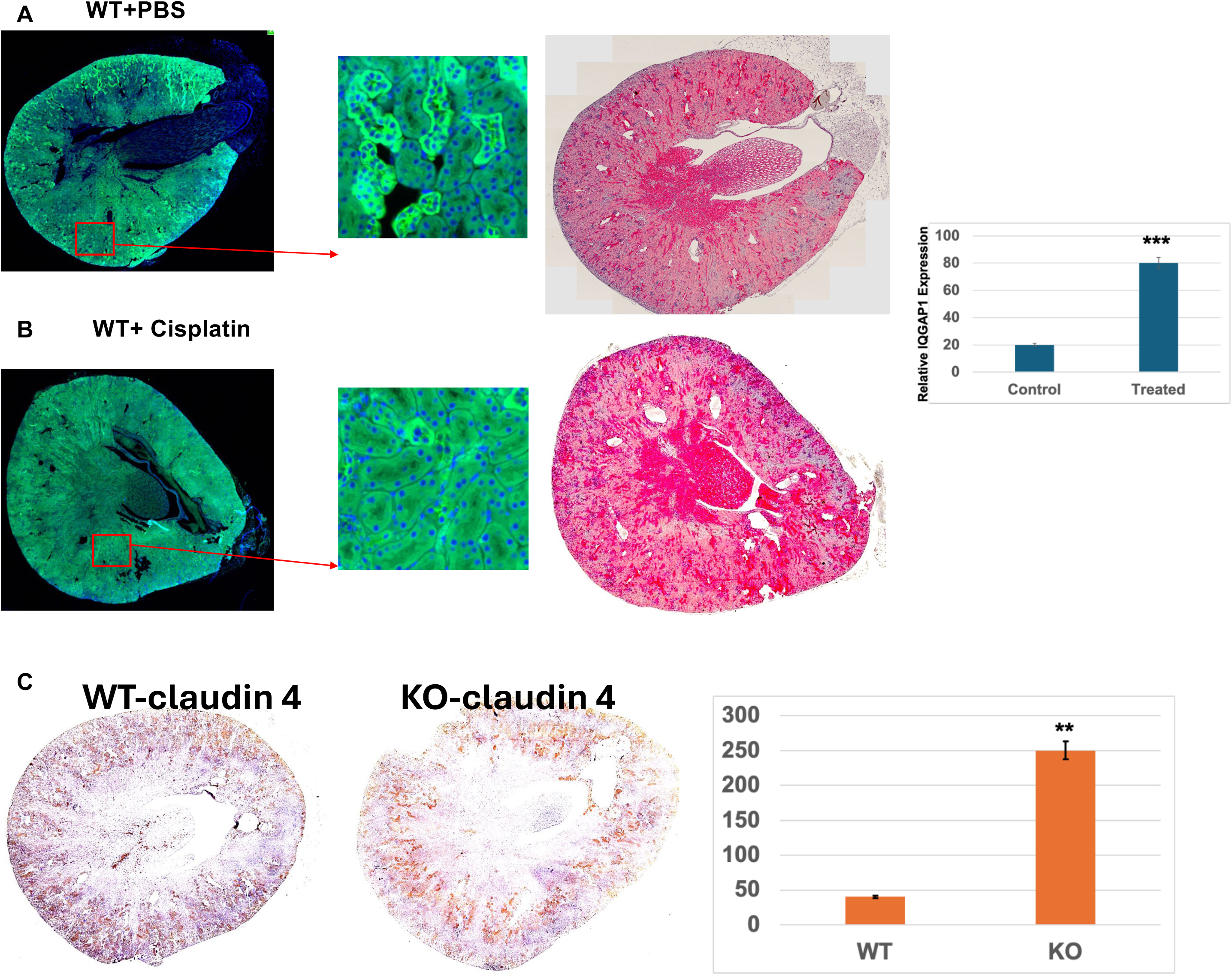
Cisplatin displaces IQGAP1 and appears to increase its level in kidney whereas loss of IQGAP1 increases claudin 4 level in the mouse kidney. **A-B**. Fluorescence IHC (green, left) and chromogen IHC (red, middle) were used to visualize IQGAP1 localization and expression level in wild type A. control (PBS-injected) or B. cisplatin-treated kidneys. The overall color signal was quantified and valued displayed in the graph (left) where the error bars are the mean ± S. D. for n = 3 for 5 random fields each. **C**. Kidneys from WT (left) and KO (right) were stained with claudin 4 antibodies in brown and the signal intensity was quantified and displayed in the graph (right) where the error bars are the mean ± S. D. for n = 3 for 5 random fields each.

